# Perineuronal nets in motor circuitry regulate the performance of learned vocalizations in songbirds

**DOI:** 10.1101/2024.05.14.593930

**Authors:** Xinghaoyun Wan, Angela S. Wang, Daria-Salina Storch, Vivian Y. Li, Jon T. Sakata

**Author notes:** **Correspondence:** <u>Corresponding author:</u> Jon T. Sakata.

## Abstract

The accurate and reliable production of learned behaviors can be important for survival and reproduction; for example, the performance of learned vocalizations (e.g., speech and birdsong) modulates the efficacy of social communication in humans and songbirds. Consequently, it is critical to understand the factors that regulate the performance of learned behaviors. Across various taxa, neural circuits that regulate motor learning are replete with perineuronal nets (PNNs), extracellular matrices that surround neurons and shape neural dynamics and plasticity. Perineuronal nets in circuits for sensory and cognitive processes have been found to affect sensory processing and behavioral plasticity. However, the function of PNNs in motor circuits remains largely unknown. Here, we analyzed the causal contribution of PNNs in motor circuitry to the performance of learned vocalizations in songbirds. Songbirds like the zebra finch are powerful models for this endeavor because the performance of their learned songs is regulated by activity within discrete and specialized circuits (i.e., song system) that are dense with PNNs. We first report that developmental increases in the density and intensity of PNNs throughout the song system [including in the motor nucleus HVC (acronym used as proper name)] are associated with developmental increases in song performance. We next discovered that enzymatically degrading PNNs in HVC acutely affected song performance. In particular, PNN degradation caused song structure to deviate from pre-surgery song due to changes in syllable sequencing and production. Collectively, our data provide compelling evidence for a causal contribution of PNNs to the performance of learned behaviors.

**SIGNIFICANCE STATEMENT:** Motor circuits are replete with perineuronal nets (PNNs) but little is known about their contribution to motor performance. Here, we analyzed how PNNs within vocal motor circuits modulate the ability of songbirds to consistently produce their learned songs. We report that developmental increases in PNN expression in vocal circuitry were associated with developmental increases in the ability to consistently perform their learned song. Moreover, degrading PNNs in the vocal motor nucleus HVC reduced the ability of adult birds to accurately produce their learned song. Our findings indicate a causal contribution of PNNs in motor circuitry to the performance of learned behaviors and, because PNNs are expressed in brain areas regulating speech, suggest that PNNs could modulate speech production in humans.

## INTRODUCTION

Many behaviors important for survival and reproduction are learned, and it is important for organisms to reliably execute these acquired behaviors. For example, spoken or signed language in humans consists of learned motor gestures and requires precise motor execution for accurate communication (Doupe and Kuhl, 1999; Horwitz et al., 2003; Long et al., 2016; Moro-Velazquez et al., 2021). Our reliance on learned motor skills such as speech and walking is underscored by the debilitating effects of neurotrauma and movement disorders on productivity and quality of life (e.g., Anderson et al., 2007; Martinez-Martin, 2017). As such, revealing how the nervous system regulates motor performance is central to our understanding of behavior and its dysfunction in disease.

Perineuronal nets (PNNs) are extracellular matrices that surround neurons within various circuits in the brain, including motor circuitry (Brückner et al., 1994; Bertolotto et al., 1996; Hausen et al., 1996; Alpár et al., 2006; Lupori et al., 2023; Wang et al., 2023). The function of PNNs has been extensively studied in sensory cortices, the hippocampus, and spinal cord, where they have been demonstrated to modulate neural activity, excitability, and plasticity (reviewed in Takesian and Hensch, 2013; Happel et al., 2014; Fawcett et al., 2019, 2022; Reichelt et al., 2019; Tansley et al., 2022). For example, PNNs in sensory circuits emerge as the sensitive period for neural plasticity closes, thereby consolidating mechanisms of sensory processing and decreasing experience-dependent plasticity, and degrading PNNs in these circuits destabilizes sensory processing and reinvigorates plasticity mechanisms (Takesian and Hensch, 2013; Banerjee et al., 2017; Fawcett et al., 2019, 2022; Tansley et al., 2022). Despite that PNNs are extensively expressed in motor circuitry, little is known about their contribution to motor performance and plasticity.

Songbirds like the zebra finch are ideal to reveal the contribution of PNNs in motor circuitry to the performance of learned behaviors. This is because songbirds learn their vocalizations (“songs”) during development (Doupe and Kuhl, 1999; Brainard and Doupe, 2013; Sakata and Woolley, 2020) and because the learning and performance of songs are regulated by discrete and specialized brain areas (“song system”) that are replete with PNNs (Balmer et al., 2009; Meyer et al., 2014; Cornez et al., 2015, 2018; Wang et al., 2023). Perineuronal net expression in songbirds resembles that in mammals in many ways; for example, similar to rodents, PNNs preferentially surround PV neurons and influence the intensity of PV expression in PV neurons (Wang et al., 2023). Previous studies have provided correlated evidence suggesting that PNNs contribute to vocal performance (Balmer et al., 2009; Cornez et al., 2015, 2018, 2020), but little is known about how PNNs actively regulate song performance. Indeed, the only study to date that has manipulated PNN expression in the song system of songbirds (canaries) documented variable effects on song performance (Cornez et al., 2021).

Here we investigated the relationship between PNN expression and vocal performance in zebra finches, focusing on the ability of birds to perform their learned songs in a consistent or reproducible manner. The ability to reliably produce acoustic features of songs a hallmark of mature song in songbirds and can influence reproductive success (Brainard and Doupe, 2000; Ölveczky et al., 2005, 2011; Woolley and Doupe, 2008; Sakata and Vehrencamp, 2012; Dhawale et al., 2017). In particular, whereas the songs of developing juvenile zebra finches are characterized by relatively high variability in the acoustic structure and sequencing of vocal elements (“syllables”), syllable structure and sequencing are highly stereotyped in the mature songs of adult finches (Brainard and Doupe, 2013; James and Sakata, 2014, 2015, 2019; Dhawale et al., 2017; Sakata and Woolley, 2020). We hypothesized that, just as PNNs in sensory circuitry consolidate sensory processing, PNNs in motor circuitry consolidate the motor commands for motor performance. We tested this hypothesis by examining developmental changes in song performance and PNN expression in the song system and by assessing how degrading PNNs in song circuitry affects the ability to perform learned songs.

## MATERIALS AND METHODS

### Animals

Thirty-five male zebra finches (*Taeniopygia guttata*) were raised in a breeding colony at McGill University. Birds were raised with both parents until 60 days post-hatch (dph) and then housed in same-sex group cages. Seven juveniles (54-70 dph) and five adults (0.8-1.2 years) were used to analyze developmental changes in song and PNN expression. Twenty-three young adults (87-133 dph) were used to reveal the effects of PNN degradation on song performance. All birds were housed on a 14:10h light:dark cycle and provided food and water *ad libitum*. All procedures were approved by the McGill University Animal Care and Use Committee in accordance with the guidelines of the Canadian Council on Animal Care.

### Song recording

Zebra finch songs were recorded in a sound-attenuating chamber (“soundbox”; TRA Acoustics, Ontario, Canada) using an omnidirectional microphone (Countryman Associates, Menlo Park, CA) positioned above the bird’s cage. Songs were recorded and digitized using Sound Analysis Pro (SAP) (2011; sampling rate: 44.1 kHz). For the analysis of developmental changes in PNN expression and song performance, songs were collected 24-48 hrs before brain collection, and for the analysis of PNN effects on song performance, songs were recorded on a daily basis starting from 24-48 hrs before surgery up to one week after surgery (when brains were collected). Recorded songs were digitally filtered at 0.3-10 kHz for off-line analysis using custom software written in MATLAB (MathWorks, Natick, MA). Birds remained individually housed throughout the experiment; therefore, all songs were undirected songs.

### Surgery and degradation of PNNs

Birds were anesthetized with an intramuscular injection of ketamine (0.03 mg/g i.m.) and midazolam (0.0015 mg/g i.m.) followed by vaporized isoflurane (0.2–3.0% in oxygen). Thereafter, birds were placed in a stereotaxic device, with their beak stabilized at a 45° angle. Bilateral craniotomy was performed (0.7-0.8 mm rostral from the caudal edge of the bifurcation of the mid-sagittal sinus, 1.8-2.0 mm lateral from the midline) and either ChABC (Sigma-Aldrich; C3667; 100 U/ml, in 0.1% BSA in PBS) or penicillinase (PEN; Sigma; P0389; 100 U/ml, in 0.1% BSA in PBS) was injected into HVC (acronym used as a proper name; 0.5 mm in depth). HVC is critical for song learning and production and regulates various aspects of song performance (reviewed in Ikeda et al., 2020; Murphy et al., 2020). Drugs were injected using a Nanoject III Programmable Nanoliter Injector (Drummond Scientific, Broomall, PA) assembled with a glass pipette at the rate of 10 nl/s (modal injection volume: 100 nl/hemisphere; range: 50-150 nl per cycle, 1-3 cycles for each hemisphere). The glass pipette was left in place for ∼2 min before retraction.

### Tissue collection and immunohistochemistry (IHC)

Birds were deeply anesthetized with isoflurane vapor, and then transcardially perfused with heparinized saline (100 IU/ml) followed by 150 ml of 4% paraformaldehyde (pH = 7.4). Brains were extracted, post-fixed overnight at 4°C in 4% paraformaldehyde, and then stored in a 30% sucrose solution at 4°C for cryoprotection. Sagittal sections were cut at 40 µm using a freezing microtome (Leica Biosystems, Wetzlar, Germany) and stored in 0.05 M Tris-buffered saline (TBS, pH = 7.6) at 4°C. For the analysis of age-dependent changes, sections from only one hemisphere were analyzed; lateralization in features measured here have not been documented (e.g., Cornez et al., 2015). Both hemispheres were analyzed to assess how bilateral PNN degradation affected song performance.

For IHC, free-floating sections were first rinsed 3X (5 min each) in 0.05 M TBS and then blocked for 30 min in 0.05 M TBS + 5.0% donkey serum + 0.1% Triton-X. Thereafter, sections were incubated for 24 h at 4°C with a mixture of a monoclonal mouse anti-chondroitin sulfate antibody (CS-56, 1:500; Sigma-Aldrich) and a polyclonal rabbit anti-PV antibody (1:2000, ab11427; Abcam) or a sheep anti-PV antibody (1:500, AF5058, Bio-Techne) in 0.05 M TBS + 0.1% Triton-X (Balmer et al., 2009; Cornez et al., 2015, 2018, 2021; Wang et al., 2023). Sections were rinsed 3X (5 min each) in 0.05 M TBS and then incubated in the dark for 2 h at room temperature with secondary antibodies: donkey anti-mouse Alexa Fluor 488 (3 µl/ml; Life Technologies) and donkey anti-rabbit Alexa Fluor 594 (5 µl/ml; Life Technologies) in 0.05 M TBS with 0.1% Triton-X. Sections were then rinsed 3X (5 min each) in 0.05 M, transferred to and stored in TBS before they were mounted on chromium-aluminum subbed slides and cover slipped (Prolong Gold Antifade; P36930; Thermo Fisher Scientific, MA, USA).

### Image acquisition and quantification

Images of PNNs and PV neurons were acquired using a Zeiss Axio Imager upright microscope and AxioCam MRm Zeiss camera (Carl Zeiss, Jena, Germany). Developmental changes in PNN and PV neuron expression in HVC, the robust nucleus of the arcopallium (RA), the lateral magnocellular nucleus of the anterior nidopallium (LMAN), and the basal ganglia nucleus Area X were investigated. For this, 3-4 40X images were acquired in the center of each brain area and used to quantify PNNs and PV neurons. For the analysis of the effects of PNN degradation on song performance, 5X images throughout HVC were taken to calculate the extent of PNN degradation and damage in HVC following ChABC or PEN infusions. To analyze how ChABC affected PV neuron density, 5-7 40X images in parts of HVC with PNN degradation were taken in ChABC birds and in similar locations in HVC of PEN birds. Exposure times were optimized for each brain area and IHC batch, and all images within each batch were taken using the same exposure times.

Neurons surrounded by PNNs (PNN neurons) and PV neurons were counted independently using Fiji (ImageJ). A PNN neuron was defined as a neuron in which ≥75% of the cell’s circumference was surrounded by continuous and bright PNN expression (Balmer et al., 2009; Wang et al., 2023). PV neurons were independently quantified in the same sections but without PNNs in the image. Thereafter, PV neurons surrounded by PNNs (PV+PNN neurons) were identified, and the percent of PV neurons that were surrounded by PNNs was computed for each bird. We could not quantify PNNs for one control (PEN) bird due to poor staining; this bird was included in behavioral analyses but excluded from analyses of PNN expression.

In addition to examining the density of PNNs, we also quantified developmental changes in the intensity of PNNs. For this analysis, 3-4 randomly chosen PNNs were outlined (regardless of whether they surrounded a PV neuron or not) per section, and the mean intensity of PNNs within the contour was quantified using ImageJ. Background intensities were subtracted from each value to yield the “PNN intensity” per PNN.

We computed the percent of HVC with PNN degradation and with damage using 5X images of PNNs and PV neurons in HVC. PV staining was used to outline and calculate the area of HVC, and the area with PNN degradation (lacking PNN staining) within HVC was computed. Insertion of pipettes in and around HVC can cause minor structural damage to HVC, and the percent of HVC with structural damage was also estimated from these sections.

Experimenters were blind to the condition of the bird when quantifying all brain images.

### Song annotation and analysis

Songs were manually labeled based on visual inspection of spectrograms following amplitude-based syllable segmentation using custom-written MATLAB software. Syllables in the songs of zebra finches are defined as individual acoustic elements separated from each other by at least 5 ms of silence, and song bouts were defined as contiguous epochs of vocalizations in which syllables are separated by <500 ms.

We analyzed song stereotypy (i.e., the similarity of vocalizations across renditions) in two ways. The songs of developing birds can be variable and difficult to annotate; in other words, because syllable structure can be highly variable, it can be difficult to accurately classify each syllable. Therefore, we first extracted the first 1.5 seconds from 20 randomly selected song bouts (“song snippets”) and then analyzed the acoustic similarities among snippets from a day of recording (i.e., acoustic similarity between different renditions of songs). These snippets started from the onset of the first introductory notes (Fig. 1), and if the 1.5-second limit fell in the middle of the syllable, we extended the snippet until the end of that syllable. The “%similarity” score between each song snippet was computed using Sound Analysis Pro (SAP) 2011 (Tchernichovski et al., 2000; Tchernichovski and Mitra, 2004; Chen et al., 2016), and the average %similarity between all snippets was calculated and used as a proxy for song stereotypy (higher %similarity indicated more stereotyped song production).

**Fig. 1.**
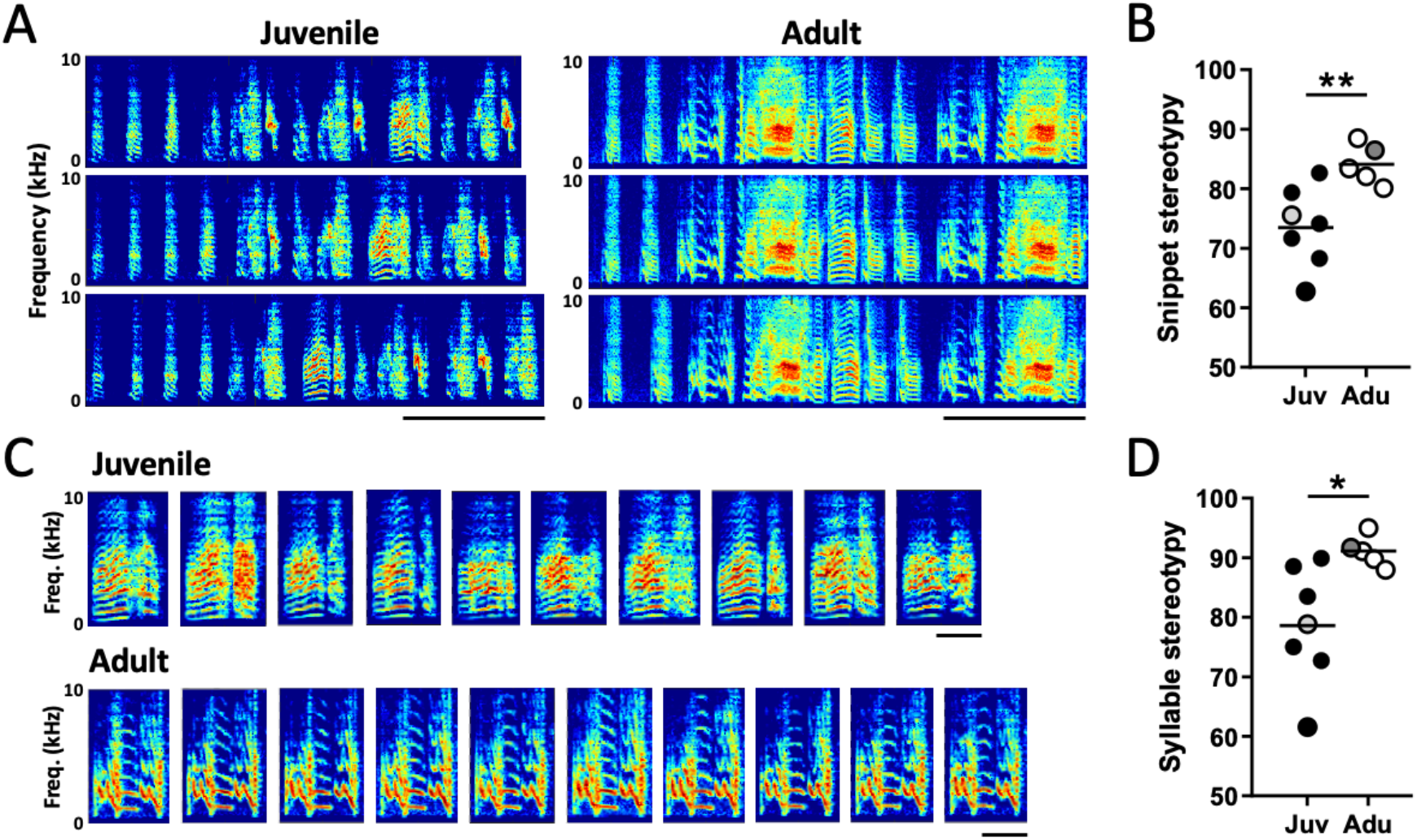
Vocal performance (stereotypy) increases with age. (A) Representative spectrograms [frequency (kHz; y-axis) vs time (ms; x-axis) with brightness representing amplitude] of three song snippets (1.5 seconds) of the songs of a juvenile (left) and adult (right) zebra finch. Scale bar: 0.5 second. (B) The acoustic stereotypy of song snippets was significantly higher for adult zebra finches than for juveniles (** p<0.01). (C) Representative spectrograms of ten renditions of one syllable type in the songs of the juvenile and adult depicted in (A). Scale bar: 0.1 second. (D) The acoustic stereotypy of individual syllables was significantly greater for adult zebra finches than for juveniles (* p<0.05). Plotted are average %similarity scores across syllable types within each bird. The gray circles in (B) and (D) indicate the data for the juvenile and adult birds depicted in (A) and (C).

Second, we analyzed the %similarity of each syllable type (e.g., “a”, “b”, “c”, etc.) that was produced ≥30 times in the 20 songs used for the snippet analysis. Specifically, after segmenting syllables within song bouts, syllables that were visually and reliably identifiable were labeled in the songs of juveniles and adults. Thereafter, 30-40 renditions of a syllable type were randomly selected; specifically, for syllables that were produced 30-40 times, all renditions were included in the analysis, and for syllables produced >40 times, 40 randomly selected renditions were analyzed. The “%similarity” score between each syllable rendition was computed (using SAP), and the average %similarity score across renditions was calculated and used as a measurement of syllable stereotypy.

Both song and syllable stereotypy measures were computed and compared across juvenile and adult male zebra finches in the analysis of developmental changes to song.

Similar analyses were conducted to reveal how degrading PNNs within HVC affected song performance. In brief, for each day (pre- and post-surgery days) in which the bird produced >20 songs, 20 randomly selected 1.5-second song snippets were extracted and song similarity across renditions within each day was computed (see above). In addition, syllables within these 20 songs were labeled, 30-40 syllable renditions were extracted, and syllable stereotypy was computed. We were able to collect 20 songs from most birds by the 2^nd^ day following surgery (15 out of 23 birds). The latency to produce 20 songs was not significantly different between ChABC- and PEN-treated birds (t(21)=0.6; p=0.5881). One ChABC bird was excluded from the analyses of syllable production because none of his syllable types were produced >30 times per day following surgery.

We also analyzed how degradation of PNNs within HVC affected the degree to which post-surgery songs resembled pre-surgery (baseline) songs (i.e., the ability of the bird to reproduce pre-surgery songs). To this end, 20 song snippets and 30-40 syllables on each post-surgery day were acoustically compared to 20 snippets and 30-40 syllables in pre-surgery songs using SAP. The average %similarity score was taken as a measure of song performance for each post-surgery day, and self-similarity scores of pre-surgery songs were used as baseline for these comparisons.

To complement the analyses of song snippet similarity following ChABC or PEN infusions, we analyzed experimental changes to syllable composition within song snippets. To this end, we labeled syllables that were reliably identifiable within the song snippets for all pre- and post-surgery days in which birds produced >20 songs and then computed the probabilities for each syllable type on each day. To quantify the degree of dissimilarity in the syllable composition of post-surgery songs from pre-surgery songs, we calculated the absolute value of the difference in syllable probabilities for each syllable type in the bird’s song (“sequence deviation”) and then summed these values across all syllable types in the bird’s repertoire. This sequence deviation value ranges from 0 (identical distributions across datasets) to 2 (no overlap in distributions across datasets).

### Statistical analyses

Generally speaking, t-tests or mixed effects models were used to analyze experimental changes to vocal performance and PNN and PV expression. Specifically, t-tests were used to analyze developmental changes to PNN density and intensity, PV neuron density, and the percent of PV neurons surrounded by PNNs (%PV+PNN) as well as to compare HVC damage between ChABC and PEN birds.

We used mixed effects models to analyze the effects of PNN degradation on PNN and PV expression, including BatchID as a random factor because brains were processed across different IHC batches (we analyzed only batches in which both ChABC and PEN birds were included in the batch; n=22 birds). Mixed effects models were also used to analyze some developmental changes in song performance and the effect of PNN degradation on song performance. For the developmental analyses, we used a mixed effects model to analyze age-dependent variation in syllable stereotypy. For this analysis, a stereotypy score (%similarity) for each syllable type in a bird’s song was used in the analysis; consequently, birdID was included as a random factor when analyzing differences between juvenile and adult zebra finches (because we analyzed multiple syllables per bird). For longitudinal analyses following PNN degradation, a full factorial model with Treatment (ChABC vs. PEN) and Time [pre-surgery day and D1-D7 (the first seven days following surgery)] and with birdID as a random variable was used to analyze changes in song performance (see above). In addition, syllable type (e.g., “a”, “b”, “c”, etc.) was also used as a random variable in the mixed effect models analyzing the syllable similarity to baseline and syllable stereotypy for ChABC and PEN birds. Planned contrasts were used to compare song performance on each post-surgery day (D1-D7) to the pre-surgery day.

Analyses were performed using JMP Pro 17 for the Mac (SAS, Cary, NC), with ?=0.05 for all omnibus tests and models.

## RESULTS

### Developmental changes in song stereotypy

We analyzed developmental changes to two different metrics of song performance, focusing on changes in the stereotypy of song performance. Developing birds produce songs that are more variable in syllable structure and sequencing, and it can be difficult to reliably identify syllables or motifs in the songs of juveniles (many analyses are based on motif structure: reviewed in Hyland Bruno and Tchernichovski, 2019). Therefore, in our first analysis, we extracted the first 1.5-second of the song starting from the first introductory note of the song bout (“song snippets”; Fig. 1A) and then computed the acoustic similarities of the snippets to each other to quantify song stereotypy (more stereotyped songs should have higher similarity scores). Song stereotypy was significantly higher for adults (n=5) than for juveniles (n=7; Fig. 1B, t(10)=3.2, p=0.0090). In addition, when we analyzed the acoustic stereotypy of individual syllables (i.e., the similarity of renditions of readily identifiable syllable types; e.g., Ölveczky et al., 2005, 2011; Fig. 1C), we observed that adult zebra finches produced significantly more stereotyped syllables than juveniles (mixed effects model: F(1,10.0)=8.2, p=0.0169; Fig. 1D). These results are consistent with previous analyses and highlight the utility of both metrics of song performance.

### Developmental changes in PNN and PV expression in the song system

Song performance is regulated by neural activity in the song system (Fig. 2A), and developmental changes in vocal stereotypy could be linked to changes in PNN expression in the song system (see also Balmer et al., 2009; Cornez et al., 2018; Fig. 2B). As such, we analyzed developmental changes in the density and intensity of PNNs in HVC, RA, LMAN, and Area X. The density of PNNs was significantly greater in adult zebra finches (n=5) in HVC (t(10)=3.1, p=0.0113) and LMAN (t(10)=2.2, p=0.0482) compared to juveniles (n=7), with similar but non-significant patterns observed in RA and Area X (Fig. 2C). The intensity of PNNs (measured in arbitrary units (a.u.); see Methods) also significantly increased over development in HVC (t(10)=2.3, p=0.0427), LMAN (t(10)=2.3, p=0.0462), and Area X (t(10)=2.6, p=0.0288; Fig. 2D). Perineuronal nets often surround PV neurons (Balmer et al., 2009; Cornez et al., 2018; Wang et al., 2023), and the percent of PV neurons surrounded by PNNs (%PV+PNN) significantly increased with age in HVC (t(10)=2.4, p=0.0348) and RA (t(10)=4.0, p=0.0025), with similar patterns in LMAN and Area X (Fig. 2E). There were no significant changes in the density of PV neurons with age in any song nucleus (Fig. 2F; p>0.14 for each).

**Fig. 2.**
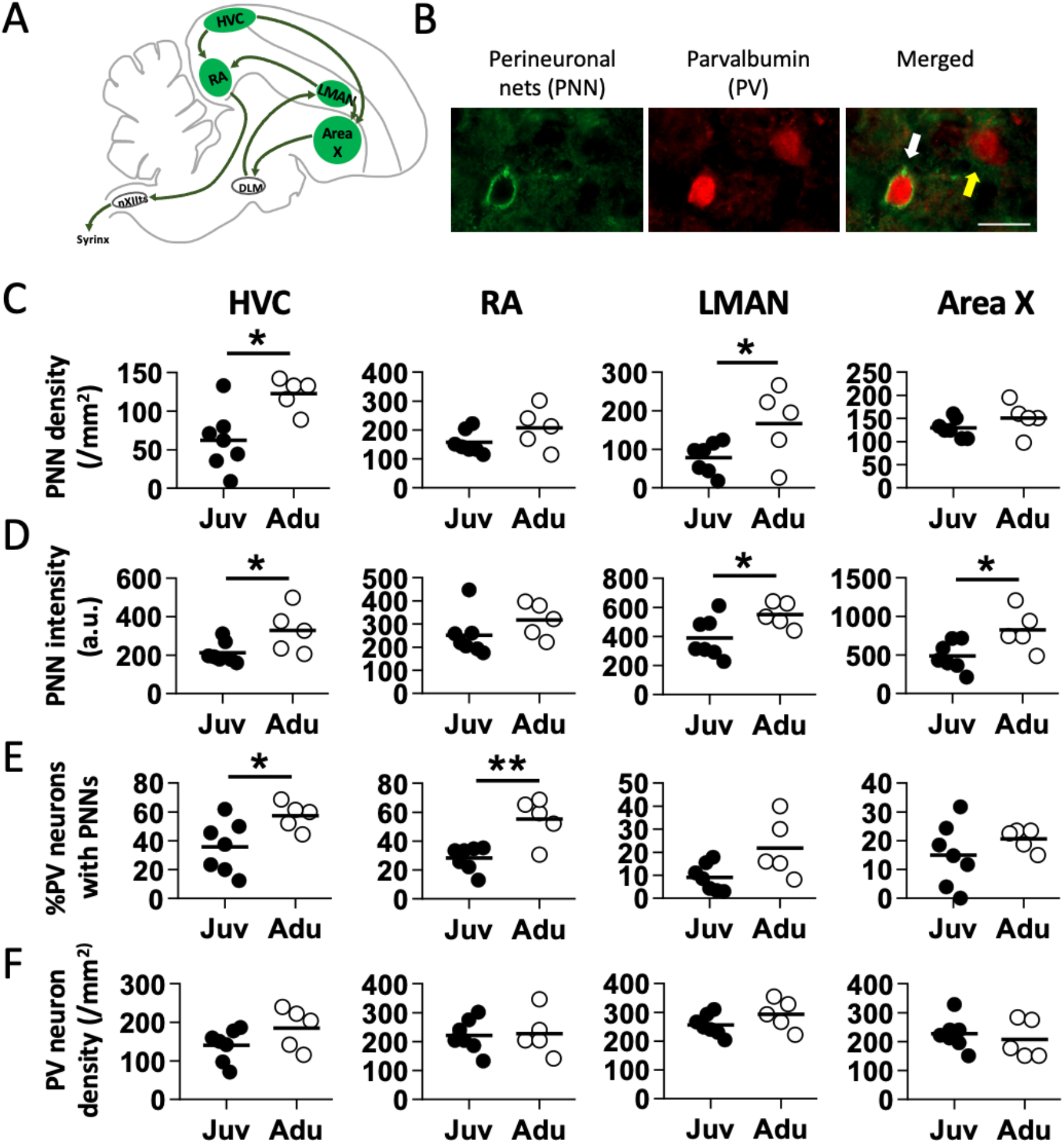
Perineuronal net expression in the song system increases with age. (A) Schematic of the song system, highlighting the forebrain areas HVC, RA, LMAN, and Area X that are replete with PNNs. (B) Representative photomicrographs illustrating a PV neuron surrounded by a PNN (white arrow) and PV neuron without PNNs (yellow arrow). Scale bar: 20 μm. Age-related changes in (C) PNN density, (D) PNN intensity, (E) the percentage of PV neurons with PNNs, and (F) PV neuron density in HVC, RA, LMAN, and Area X (* p<0.05, ** p<0.01).

### Effects of PNN degradation on the performance of learned song

Given the consistent associations between age-dependent changes in song performance and in PNN expression in HVC, we investigated how manipulations of PNN expression in HVC affected song performance. HVC is critical for the production of learned songs and regulates various aspects of song performance (reviewed in Murphy et al., 2020). In particular, we tested the hypothesis that degrading PNNs in HVC would decrease song performance by (a) decreasing the similarity of post-surgery song to pre-surgery (baseline) songs and/or (b) decreasing the stereotypy of song performance.

Stereotaxic injection of ChABC into HVC (n=13) caused conspicuous PNN degradation whereas injection of PEN into HVC (n=10) left PNNs largely intact (Fig. 3A). Not surprisingly, there was significantly greater PNN degradation within the HVC of ChABC birds than of PEN birds (Fig. 3B; F(1,18.1)=112.0, p<0.0001). There was no significant difference in the density of PV neurons between the two groups (Fig. 3C; F(1,17.7)=1.2, p=0.2967), consistent with previous reports that enzymatic degradation of PNNs does not affect neuron numbers (Cabungcal et al., 2013; Yamada and Jinno, 2013; Yamada et al., 2015). Additionally, the degree of damage to HVC was not significantly different between ChABC and PEN birds (F(1,20.3)=2.2, p=0.1572).

**Fig. 3.**
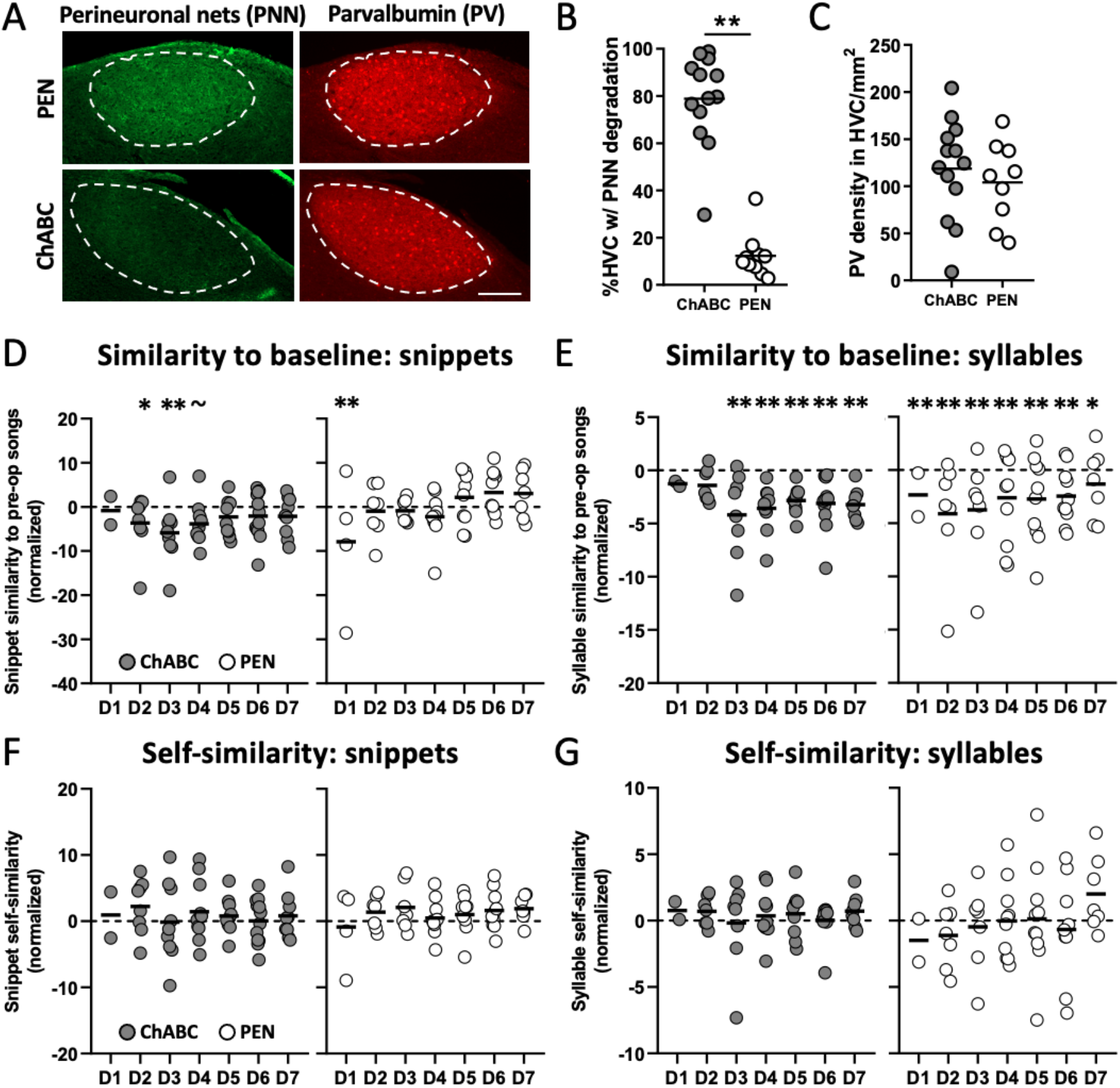
Degrading PNNs in HVC affected the ability to perform learned songs but did not affect vocal stereotypy. (A) Representative photomicrographs illustrating PNN (green) degradation caused by ChABC (bottom row) and intact PNNs in control birds that received the injection of PEN (top row). The white dashed line outlines HVC in each panel (based on PV expression). Scale bar: 200 μm. (B) The percentage of HVC with PNN degradation was significantly higher in ChABC birds than in PEN birds (** p<0.01). (C) There was no significant difference in PV neuron density in HVC between ChABC and PEN birds (p>0.25). (D-E) The similarity of post-surgery song snippets (D) and individual syllables (E) to pre-surgery songs or syllables (** p<0.01, * p<0.05, ∼ p<0.06). (F-G) The acoustic stereotypy of song snippets (F) and individual syllables (G) was not affected by PNN degradation (p>0.05).

To assess the effects of PNN degradation on song performance, we collected and compared pre-surgery songs (“baseline”; recorded 1-2 days before surgery) and post-surgery songs of ChABC- and PEN-treated birds. In general, we did not visually observe dramatic changes to song structure following either ChABC or PEN infusions into HVC. However, we analyzed how infusions affected the acoustic similarity of post-surgery song snippets to pre-surgery (“baseline”) song snippets (n=20 snippets per day; see Methods). We also computed the similarity of baseline song snippets to themselves to represent baseline levels of song similarity (“baseline similarity”). A full-factorial model [Treatment (ChABC vs. PEN) x Time (baseline and post-surgery days 1-7 (D1-D7))] indicated a trend for a Treatment x Time interaction (mixed effects model: F(7,103.6)=2.0, p=0.0593; Fig. 3D). This was driven by the fact that the patterns and magnitudes of changes in song performance (i.e., similarity to baseline song) differed between ChABC and PEN birds. Following ChABC infusions, similarity to baseline song was significantly decreased on post-surgery days 2 and 3 (p<0.05), with a strong trend for post-surgery day 4 (p<0.06). There was also a transient decrease in song performance following PEN infusions but the decrease was significant only on post-surgery day 1 (p<0.01). In summary, infusions that degrade PNNs in HVC lead to a more prolonged song deterioration than control infusions.

We next sought to understand how variation in syllable structure or sequencing could account for the differential decrease in song performance in ChABC birds. We first examined how ChABC or PEN infusion into HVC affected the acoustic similarities of post-surgery syllables to pre-surgery (“baseline”) syllables. Interestingly, syllable structure changed in different ways following ChABC or PEN infusions into HVC (Treatment x Time interaction: F(7,368)=2.0, p=0.0483; Fig. 3E). Following ChABC infusions, syllable structure was different on post-surgery days 3-7 compared to baseline (p<0.05 for each planned contrast), with a trend for post-surgery day 2 (p=0.0527). On the other hand, syllable structure was different on post-surgery days 1-7 for PEN birds (p<0.05 for each). All in all, these data underscore comparable changes to syllable performance for ChABC and PEN birds and suggest that variation in the effects of ChABC and PEN on song performance (Fig. 3D) cannot be readily explained by differential effects of ChABC and PEN on syllable performance (Fig. 3E).

We next investigated the degree to which differential changes to syllable sequencing could contribute to the differential change in song similarity. For example, it is possible that song similarity decreased in ChABC but not PEN (control) birds on post-surgery days 2-4 because syllable composition within extracted snippets (see Methods) changed following surgery for ChABC but not for PEN birds. Phrased differently, it is possible that degradation of PNNs in HVC affected syllable sequencing, in particular sequencing at the beginning of song. To test this, we labeled all syllables in the extracted song snippets and then created a histogram of syllable prevalence (probably of each syllable type) for each pre- and post-surgery day. We then computed the sequence deviations (see Methods) between the songs from each post-surgery day to pre-surgery songs (larger values indicate histograms that are more different from each other). Examples of syllable compositions within pre-surgery and post-surgery days for a bird that received ChABC infusions bilaterally into HVC and for a bird that received PEN infusions bilaterally into HVC are provided in Fig. 4A,B. Syllable compositions within snippets changed more for the ChABC bird than for the PEN bird. When sequence deviations were analyzed across all birds, it was found that the magnitude of sequence deviations was significantly greater for ChABC birds than for PEN birds (F(1,25.1)=5.1, p=0.0336; Fig. 4C). While sequence deviations from baseline songs were consistently greater for ChABC birds than for PEN birds following surgery, the difference in the magnitude of sequence deviation was greatest on post-surgery day 3. This temporal pattern of sequence variation resembles that of variation in song similarity, suggesting that changes in syllable sequencing (i.e., syllable composition within the extracted snippets of song) account for change in song performance.

**Fig. 4.**
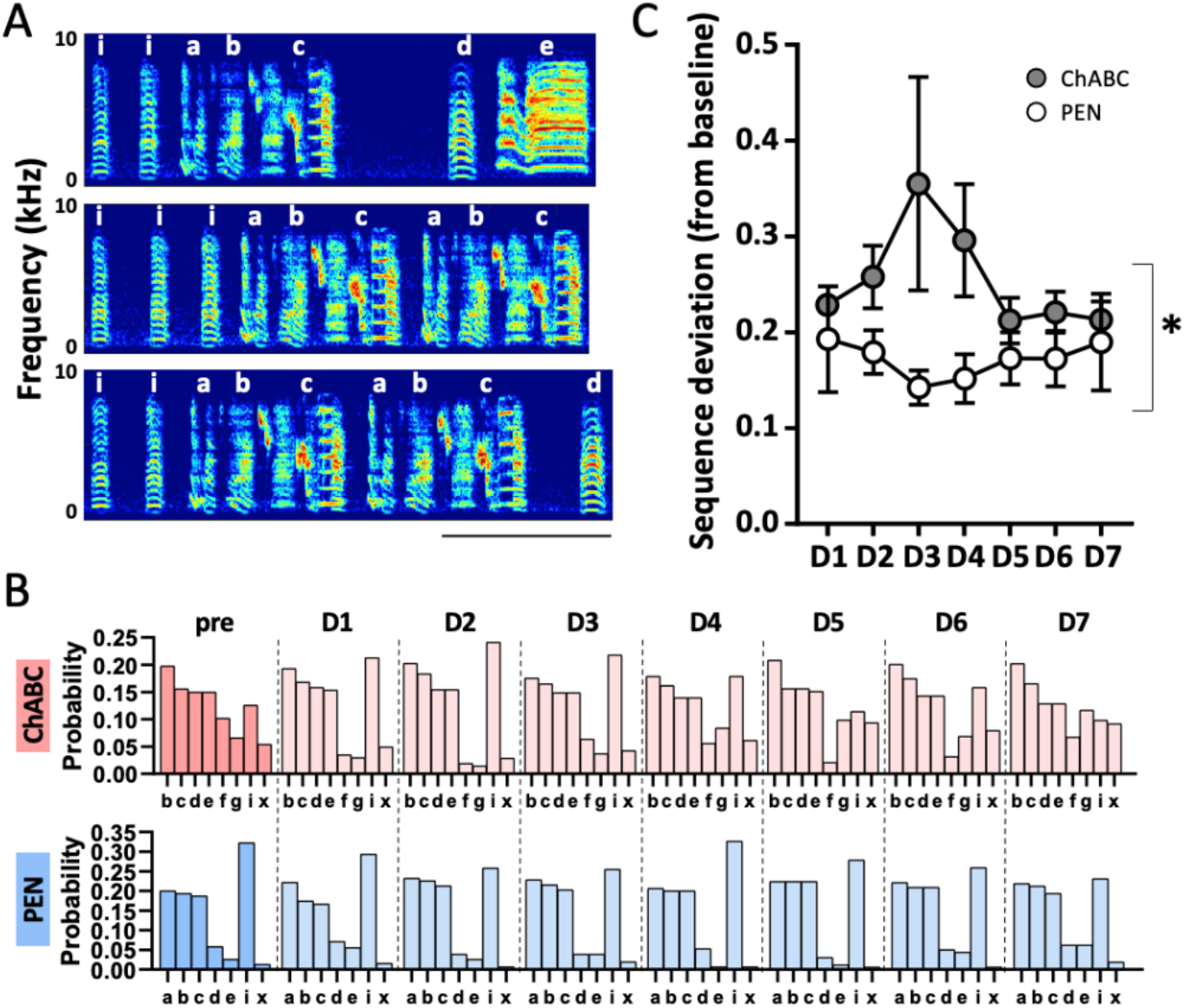
ChABC infusions cause significant changes to syllable sequencing. (A) Representative spectrograms of three song snippets (1.5 seconds) with syllables (white letters) from a PEN bird (pre-surgery). Scale bar: 0.5 second. (B) Histograms of syllable composition (proportions) within pre-surgery and post-surgery days for a ChABC bird (top row; pink) and a PEN bird (bottom row; blue). (C) The magnitude of sequence deviations from baseline song was significantly higher in ChABC birds than in PEN birds (*p<0.05; see text).

Song snippets (from the beginning of song) consist of introductory notes and other song syllables. Introductory notes are repeated at the beginning of song and changes in the number of repetitions of introductory notes could underlie sequence changes following PNN degradation. However, there was no significant difference in the proportion of syllables within song snippets that were introductory notes between ChABC and PEN birds (F(1,21.5)=1.0, p=0.4142), nor was there a significant interaction between Treatment and Time (F(7,97.6)=1.6, p=0.1338). This suggests that changes in the repetition of introductory notes did not account for the differential change in syllable sequencing following PNN degradation.

In addition to analyzing how PNN degradation affected the ability of birds to reproduce their pre-surgery songs, we also analyzed the degree to which ChABC vs. PEN infusions differentially affected the stereotypy of song performance. Perineuronal net degradation did not differentially affect either song stereotypy (Treatment x Time interaction: F(7,103)=1.1, p=0.3875) or syllable stereotypy (Treatment x Time interaction: F(7,367)=1.2, p=0.3070 ; Fig. 3F, 3G).

## DISCUSSION

The contributions of PNNs to sensory processing and cognition have been extensively studied (reviewed in Fawcett et al., 2019, 2022). However, despite that PNNs are abundant throughout motor circuitry in various vertebrate taxa (Balmer et al., 2009; Cornez et al., 2015, 2017; Ueno et al., 2019; Carulli et al., 2020; Dzyubenko et al., 2023; Lupori et al., 2023; Wang et al., 2023), little is known about how PNNs regulate motor performance. Because PNNs in sensory circuits stabilize experience-dependent sensory processing (Pizzorusso et al., 2002; McRae et al., 2007; Gogolla et al., 2009), we tested the hypothesis that PNNs in motor circuits serve to stabilize the motor commands for learned behaviors. We specifically investigated how the expression of PNNs in the vocal motor circuit of songbirds (“song system”) correlates with and actively regulates the ability of birds to consistently produce their learned songs. We report that developmental increases in PNN expression in various areas of the song system were associated with developmental increases in the ability to consistently produce learned songs and that degradation of PNNs in the vocal motor nucleus HVC reduces the ability of adult birds to accurately produce their learned song.

Consistent with previous investigations, we report that adult zebra finches produced more stereotyped songs than juveniles (Tchernichovski et al., 2001; Ölveczky et al., 2011) and that PNN expression is higher in adults than in juveniles in three song nuclei: HVC, RA, and LMAN (Balmer et al., 2009; Cornez et al., 2018). Although these aspects of vocal performance and PNN expression have been analyzed in separate studies, no study to date has analyzed them in the same population of zebra finches (but see Cornez et al., 2020 in canaries). There was some variation across areas in the song system in the degree to which PNN expression changed over development, however PNN intensity and density and the percent of PV neurons surrounded by PNNs all increased over development in HVC. The degree to which changes in PNN intensity and density are mechanistically linked is unclear, but both could be due to an increase in the synthesis of PNN components or a decrease in cellular processes that degrade PNNs. Moreover, the coordinated changes in PNN expression and vocal performance suggest that PNNs could be involved in the ability of birds to consistently produce their learned song.

Degrading PNNs in HVC decreased the ability of birds to reproduce their learned (baseline) songs, supporting findings from our developmental analysis of song performance and PNN expression. We propose that PNN degradation affected song performance primarily by altering syllable sequencing. This is because birds with ChABC infusions into HVC demonstrated significantly greater sequence changes than birds with PEN (control) infusions and because changes in the acoustic structure of syllables (and therefore decreases in the similarity of post-surgery syllables to baseline syllables) were comparable between ChABC and PEN birds. This decrease in the ability to produce learned song qualitatively resembles how manipulations of HVC activity modulate song performance (e.g., Kosche et al., 2015; Isola et al., 2020), in particular syllable sequencing (e.g., Vu et al., 1994; Ashmore et al., 2005; Wang et al., 2008; Kosche et al., 2015; Picardo et al., 2016; Isola et al., 2020; Jaffe and Brainard, 2020). For example, manipulations of cholinergic tone (e.g., infusions of carbachol) and of inhibition in HVC affects various aspects of syllable sequencing in a related songbird, the Bengalese finch (Isola et al., 2020; Jaffe and Brainard, 2020). This suggests that PNN degradation might lead to changes in inhibition or cholinergic tone to affect syllable sequencing.

Because neural dynamics in HVC regulate song performance (reviewed in Murphy et al., 2020), our data suggest that PNNs within motor circuitry shape neural dynamics and, thus, the motor commands for song. In various parts of the rodent brain, PNNs have been found to affect neural dynamics (Liu et al., 2013; Lensjø et al., 2017; Tewari et al., 2018; Carceller et al., 2020; Carulli et al., 2020). For example, degrading PNNs in the visual, motor, and somatosensory cortices reduces spiking activity of inhibitory neurons (Lensjø et al., 2017; Tewari et al., 2018), and degrading PNNs in the deep cerebellar nuclei (DCN) decreases the spontaneous activity of DCN neurons (Carulli et al., 2020). In addition, PNNs preferentially surround parvalbumin (PV) neurons in songbirds and rodents (Balmer et al., 2009; Cornez et al., 2018; Reichelt et al., 2019; Wang et al., 2023), and PV neurons are key regulators of neural dynamics (Cardin et al., 2009; Sohal et al., 2009; Albéri et al., 2013; Hu et al., 2014; Xue et al., 2014; Reichelt et al., 2019; Nocon et al., 2023). Degradation of PNNs has been found to affect the expression of PV within PV neurons (Beurdeley et al., 2012; Yamada et al., 2015; Hou et al., 2017; Rowlands et al., 2018; Wang et al., 2023) and PV neuron activity (Balmer, 2016; Carceller et al., 2020); consequently, changes to vocal motor performance following PNN degradation in HVC could be linked to changes in PV neurons. Although we did not report significant changes to the abundance of PV neurons (see also Cornez et al., 2021), PNN degradation could lead to other changes in PV neuron function that could regulate circuit dynamics within HVC and drive changes to vocal sequencing.

The current findings complement and help interpret an existing examination of PNN contributions to the performance of learned songs (Cornez et al., 2021). That study analyzed how degrading PNNs in HVC affected the performance of canary song from 1-10 weeks after PNN degradation. The authors reported some suggestive and preliminary evidence of PNN contributions to adult song production; for example, although the effects varied across experiments, PNN degradation could lead to changes in syllable repertoire and amplitude. The relatively small and inconsistent effects across the experiments in their study could be due to variability in the timing of song recording and analysis, the reproductive state and age of the canaries, and the degree of PNN degradation and recovery. We discovered changes in the performance of zebra finch song within days of PNN removal, thereby highlighting relatively acute contributions of PNNs to song performance. However, many aspects of song performance recovered to pre-surgery levels within a week, suggesting that more consistent effects on canary song might have been observed if researchers were able to analyze songs produced within days following surgery.

Taken together, by revealing the contribution of PNNs to the performance of learned vocal motor behaviors, these experiments expand our understanding of the general function of PNNs and complement studies of PNNs function in sensory and cognitive circuitry. Given that PNNs are replete in the motor cortex, basal ganglia, and cerebellum of mammals (e.g., Lee et al., 2012; Richard et al., 2018; Briones et al., 2022), our data suggest that PNNs in motor circuitry modulate behavioral learning and performance. Of particular interest is that PNNs are expressed in brain areas underlying speech performance, including the sequencing of speech syllables (Bohland and Guenther, 2006; Virginito et al., 2009; Rong et al., 2018), and we propose that PNNs in these areas modulate speech performance. Finally, given the role of PNNs in restricting neural plasticity (Pizzorusso et al., 2002; Beurdeley et al., 2012; Takesian and Hensch, 2013; Reichelt et al., 2019) and the roles of the song system in song acquisition and development, our experiments motivate investigations into the role of PNNs in the sensory and sensorimotor learning of song.

## Abbreviated title

PNNs in motor circuitry regulate vocal performance: 

## Acknowledgements

We thank F. Bendall, O. Ruge, A. Meilayi, and Y.X. Chen for their help with data processing. Thank you also to S.C. Woolley, E. de Villers-Sidani, and A. Watt for input on data analysis and presentation. Research was supported by the Natural Sciences and Engineering Research Council (NSERC; #05016 to J.T.S.), the Fonds de recherche du Québec–Nature et technologies (FRQNT) grant to J.T.S. (PR-284884), and funds from the Centre for Research for Brain, Language, and Music (CRBLM) to J.T.S.

